# Engineering a 3D Lung Co-culture Platform to Model Epithelial–Fibroblast Interactions in Pulmonary Fibrosis

**DOI:** 10.64898/2026.05.07.723587

**Authors:** Mejalaa Mega Jayaseelan, Aleksander Skardal, Landon W. Locke, Megan N. Ballinger

## Abstract

Idiopathic pulmonary fibrosis (IPF) is a fatal interstitial lung disease (ILD) characterized by progressive fibrosis, irreversible loss of lung elasticity, and chronic respiratory failure, with a mean survival of 3–5 years. The disease is believed to result from repeated alveolar epithelial injury that sustains transforming growth factor-beta (TGF-β) signaling, driving fibroblast-to-myofibroblast differentiation and excessive collagen deposition. Although current IPF models—including animal studies, 2D cultures, and basic 3D systems—have enhanced understanding of disease mechanisms, they inadequately replicate epithelial–fibroblast interactions, extracellular matrix (ECM) remodeling, and epithelial barrier dysfunction. To address this limitation, we engineered a 3D lung co-culture model that simulates the physiological epithelial–fibroblast crosstalk and ECM remodeling characteristic of IPF. Our model embeds fibroblasts within a collagen–hyaluronic acid matrix overlaid with an epithelial monolayer cultured at an air–liquid interface. Basolateral TGF-β exposure generated a profibrotic microenvironment that weakened epithelial barrier integrity and drove myofibroblast differentiation marked by elevated α-SMA and vimentin. Elevated pro-inflammatory cytokine secretion and increased collagen disorganization further demonstrated active fibrogenesis. Together, these features show that our model captures key early events in IPF pathogenesis and offers a versatile platform for next-generation lung-on-a-chip studies in fibrotic disease.

## 1. Introduction

The lung is a highly specialized organ optimized for gas exchange, oxygen uptake and carbon dioxide removal. Its structure consists of a branching airway network that ends in vascularized alveolar sacs, which enables rapid diffusion. Because the lung is continuously exposed to pathogens, particulates, and environmental toxins, it has evolved a wound healing response involving epithelial regeneration, immune activation, and matrix remodeling. Idiopathic pulmonary fibrosis is a chronic, progressive interstitial lung disease marked by excessive extracellular matrix (ECM) deposition, fibroblast activation, and irreversible destruction of alveolar architecture. With a median survival time of 3–5 years after diagnosis, it remains one of the most aggressive forms of interstitial lung disease with limited therapeutic options (Golchin et al., 2025; Olson et al., 2018; Quinn et al., 2019). Repeated epithelial microinjury is thought to trigger aberrant wound healing responses driven by transforming growth factor-beta (TGF-β). Normally secreted in a latent form, TGF-β is activated by integrins such as αvβ6 during epithelial damage (Frangogiannis, 2020). Active TGF-β promotes epithelial-to-mesenchymal transition (EMT), increases barrier permeability, suppresses epithelial cell proliferation, and induces fibroblast differentiation into matrix-remodeling myofibroblasts (Deng et al., 2024; Koukourakis et al., 2017). The resulting collagen-rich matrix drives tissue stiffening, fiber bundling, and architectural distortion, while persistent TGF-β signaling sustains inflammation, myofibroblast accumulation, and non-resolving fibrosis. Despite advances in understanding its molecular drivers, current FDA-approved treatments only slow disease progression and do not halt or reverse existing fibrosis, underscoring the urgent need for new therapeutic targets and more predictive disease models.

Understanding the molecular drivers of fibrosis requires experimental models that faithfully recapitulate its multicellular complexity. Animal models, such as the bleomycin-induced lung fibrosis model, have been essential for studying pulmonary fibrosis and testing therapies, but even refined multi-hit approaches still fail to fully reproduce the chronic, progressive pathology observed in humans (Chen et al., 2025; Tashiro et al., 2017). To overcome the limitations of animal models, a range of *in vitro* systems have been developed to dissect the molecular mechanisms of fibrosis. The most basic are two-dimensional monocultures of fibroblasts or alveolar epithelial cells and although useful for studying processes such as fibroblast-to-myofibroblast differentiation and EMT, they lack cellular diversity, ECM complexity, and mechanical cues that shape fibrotic signaling. Because 2D systems cannot reproduce the cell–cell and cell–matrix interactions or the dynamic mechanical forces that shape fibrotic signaling, they fall short of modeling key aspects of fibrosis pathogenesis (Goldmann et al., 2018; Kolanko et al., 2024).

Three-dimensional model systems offer a more physiological relevant alternative, often using hydrogel-based scaffolds that better mimic native lung architecture and enable investigation of cell migration, traction dynamics, and integrin adhesions (Leonard-Duke et al., 2020; Phogat et al., 2023). Accordingly, we engineered a three-dimensional model using a collagen–hyaluronic acid hydrogel (Mazzocchi et al., 2018), a matrix combination that integrates collagen’s structural and integrin-binding cues with hyaluronic acid’s capacity to regulate hydration, viscoelasticity, and cell signaling, thereby recreating key biochemical and mechanical features of the fibrotic lung microenvironment. This platform provides a controlled, biologically meaningful environment for probing epithelial–mesenchymal crosstalk, dissecting fibrogenic mechanisms, and evaluating interventions that modulate epithelial injury, fibroblast activation, and matrix remodeling.

## 2. Methods

### 2.1 Cell Culture

A549 human lung adenocarcinoma epithelial cells (ATCC, Cat. #CCL-185) were cultured on standard 60 mm tissue culture dishes in Dulbecco’s Modified Eagle Medium (DMEM), which was supplemented with 10% fetal bovine serum (FBS), L-glutamate, and penicillin-streptomycin. Cultures were maintained at 37°C in a humidified incubator with 5% CO_2_. When cells reached approximately 70–80% confluence, monolayers were rinsed with phosphate-buffered saline (PBS) and detached by incubation with 0.05% trypsin–EDTA solution. Detached cells were centrifuged, resuspended in fresh complete DMEM, and then re-plated at a density of 1 × 10^6^ cells per 60 mm dish for subsequent expansions and experimental use. The media was replenished every 2 days.

Normal human lung fibroblasts (NHLF; Lonza, Cat. #2512) were cultured in T75 tissue culture flasks containing complete Fibroblast Basal Medium (FBM) supplemented as per manufacturer guidelines (including insulin, rhFGF-B, GA-1000, and FBS). NHLF cultures were maintained under identical incubator conditions as the A549 cells. Only NHLFs between passage number 5 and 9 were used for all experiments to ensure consistency and avoid passage-related phenotypic drift. Upon reaching 70–90% confluence, NHLF monolayers were passaged by trypsinization, counted for viability, and re-plated at a density of 1 × 10^6^ cells per T75 flask. The media was replenished every other day.

### 2.2 Hydrogel Construct Biofabrication

Hydrogels were formulated using methacrylated collagen type I (PhotoCol; Advanced Biomatrix, Cat# 5198), thiol-modified hyaluronan with heparin (Heprasil; Advanced Biomatrix, Cat# GS215), and the photoinitiator Irgacure 2959 (2-Hydroxy-4′-(2-hydroxyethoxy)-2-methylpropiophenone; Sigma Aldrich, Cat# 410896) (Mazzocchi et al., 2018). Components were combined with a neutralization solution provided by the supplier. NHLFs were first resuspended at 3 × 10^7^ cells/mL and centrifuged to form a pellet, after which the hydrogel components were added sequentially directly onto the pellet in the Eppendorf tube. The mixture was gently pipetted to ensure homogeneous cell distribution throughout the pre-polymer solution.

For monoculture constructs, 10 µL of the NHLF-hydrogel suspension was dispensed centrally into each well of a 48-well tissue culture plate and crosslinked with a BlueWave RediCure 365 UV Lamp (365 nm) for 2 seconds. Following polymerization, 300 µL of complete FBM was added to each well and replaced every other day. TGF-β treatment groups received recombinant human TGF-β1 (10 ng/mL, Peprotech) diluted in FBM, added directly to the medium. The treatment duration was 4 days, with media replenished on day 2, after which constructs were fixed in 4% paraformaldehyde (PFA) for downstream analysis.

### 2.3 3D Transwell Models

For co-culture transwell constructs, NHLFs were pelleted at a density of 6 × 10^7^ cells/mL and suspended in the hydrogel mixture. A total of 20 µL of the gel-cell suspension was carefully pipetted into each 6.5 mm diameter polyester transwell insert (0.4 µm pore, Corning) to ensure full coverage of the permeable membrane. Gels were crosslinked with UV light 2 seconds. After polymerization, A549 cells were seeded directly onto the hydrogel surface at a density of 2.5 × 10^5^ cells/insert in 100 µL of complete DMEM. After 24 hours of submerged culture, the apical medium was removed to initiate air–liquid interface (ALI) conditions. Co-cultures were maintained for 5 days at ALI, after which TGF-β treatment (10 ng/mL) was initiated from the basolateral chamber for an additional 4 days. Media was refreshed once during the treatment period. At the conclusion of the experiment, constructs were excised from transwells and fixed in 4% PFA for further analysis.

### 2.4 TEER

Barrier integrity of the A549 epithelial layer was assessed by transepithelial electrical resistance (TEER) measurements using a commercially available TEER measurement system as we have previously described (DePalma et al., 2025). One electrode was placed in the apical chamber of the transwell and the other in the corresponding basolateral chamber. Two readings were recorded per insert, with three biological replicates per condition. Measurements were averaged to obtain a representative TEER value for each time point. TEER was monitored daily over a 5-day period to evaluate epithelial barrier formation and was continued for an additional 4 days during TGF-β treatment to assess barrier disruption.

### 2.5 Picrosirius Red

Constructs were fixed in 4% paraformaldehyde, dehydrated through graded ethanol series, and embedded in paraffin wax. Paraffin-embedded constructs were sectioned at 6 μm thickness and mounted on glass slides. Deparaffinized sections were stained with Picrosirius Red using a commercial staining kit (Abcam, Cat# ab150681), following the manufacturer’s instructions. Slides were mounted and imaged under polarized light microscopy to visualize collagen fiber deposition and organization. Regions of interest (ROIs) were selected from each sample (n ≥ 6 ROIs, field size 100 × 100 μm), and quantitative analysis of fiber architecture was performed using CT-FIRE software (Bredfeldt et al., 2014). To assess collagen fiber orientation, individual fiber angle data obtained from CT-FIRE were post-processed using a custom MATLAB script to generate rose plots.

### 2.6 IF Staining

Fixed samples were permeabilized and blocked in a solution containing 5% normal goat serum, 0.1% Triton X-100, and 0.1% sodium azide. Primary antibodies included: α-SMA (ThermoFisher Cat. #14-9760-82), vimentin (SantaCruze Cat. #sc-373717), ZO-1(ThermoFisher Cat. #339188), phalloidin (Thermo Fisher Cat. #A12379) for F-actin, and nuclei were counterstained using DAPI (Thermo Fisher Cat. #R37606). After overnight incubation at 4 °C, samples were washed and incubated with species-appropriate fluorescent secondary antibodies. Following final washes, constructs were imaged using a fluorescence microscope. α-SMA and vimentin fluorescence were quantified in ImageJ/Fiji by measuring background-subtracted integrated fluorescence intensity within each image, and values were normalized to the number of nuclei per image to report fluorescence per cell. ZO-1 junction organization was quantified in ImageJ/Fiji by thresholding the ZO-1 channel to generate binary junction masks, followed by skeletonization and analysis using the Analyze Skeleton (2D/3D) plugin in Fiji to extract total branch length and junction-related features per image.

### 2.7 ELISAs

Cytokine secretion was measured in basolateral media collected from the transwell co-culture system at two time points: day 2 (±TGF-β) and day 4 (±TGF-β). Samples were collected, spun down to remove debris, and stored at −80 °C until analysis. Prior to use, samples were thawed on ice and diluted as needed to remain within the standard curve range. Human IL-6 (Abcam, Cat# ab178013), IL-1β (Cat# ab214025), and IL-8 (Cat# ab214030) levels were quantified using ELISA kits per manufacturer protocols. For each experimental condition, supernatants from three independent biological replicates were analyzed, and each ELISA was performed in technical duplicate. Absorbance was recorded using the Varioskan Lux Microplate Reader (Thermo Fisher), and cytokine concentrations were interpolated from standard curves generated by prism from recombinant protein standards.

### 2.8 Statistical analysis

All data are presented as mean ± SEM, with individual data points representing independent biological replicates (n ≥ 3 per condition). Ordinary one-way ANOVA with Tukey’s post hoc multiple-comparisons test was used to analyze group differences for Figures 1–3. Cytokine measurements in Figure 4 were analyzed by two-way ANOVA (factors: time and TGF-β treatment) with Tukey’s post hoc test for selected pairwise comparisons. Statistical analyses and graph generation were performed using GraphPad Prism (version 11.0.0 (93), GraphPad Software, San Diego, CA). A p-value < 0.05 was considered statistically significant.

**Figure 1.**
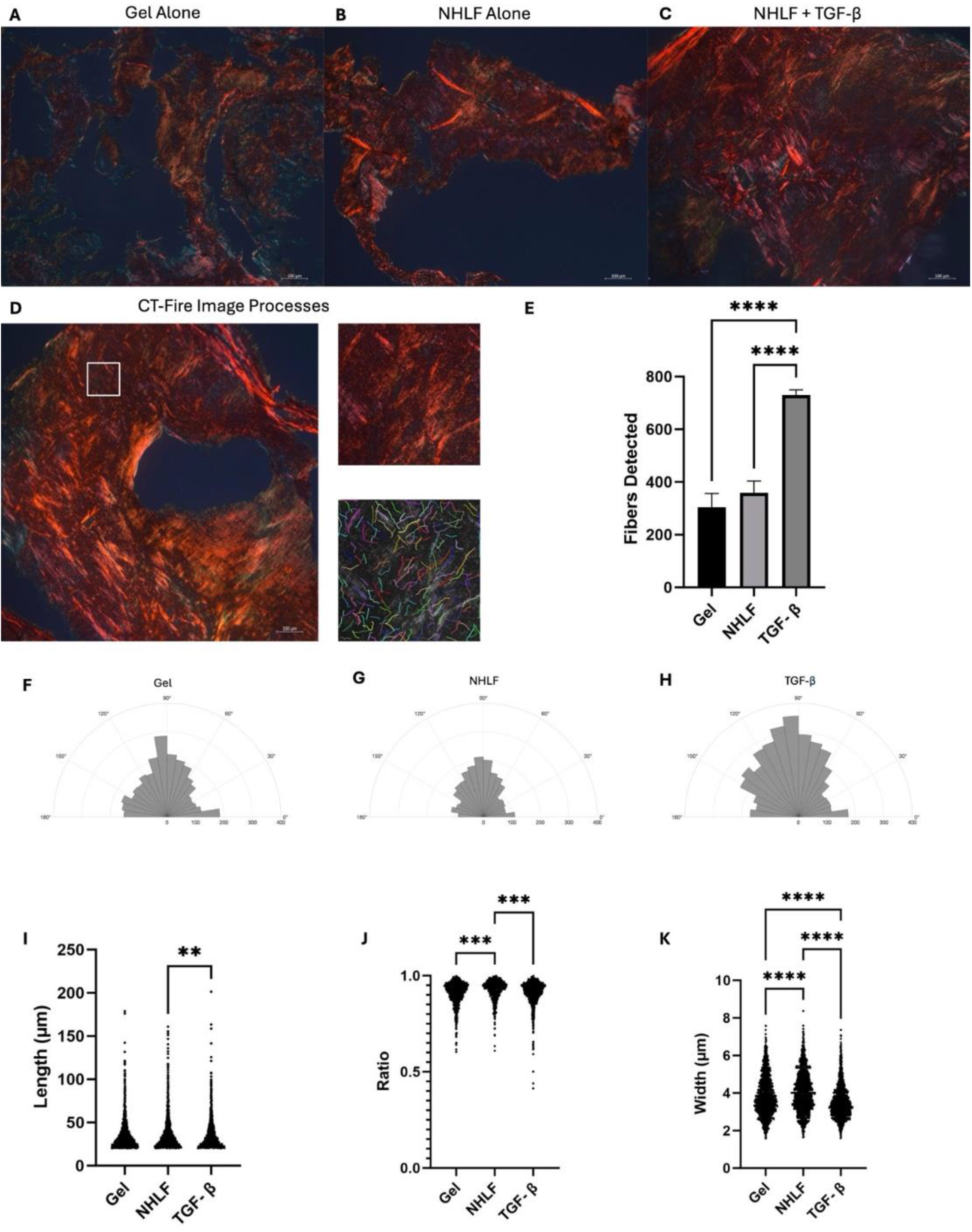
TGF-β promotes collagen remodeling and fiber alignment in 3D Hydrogel Constructs. (A) Picrosirius red stained gel only construct (B) Picrosirius red stained NHLF laden construct (C) Picrosirius red stained NHLF exposed to TGF-β sample (D) Representative CT-Fire image processing workflow, including region-of-interest selection, and automated single-fiber tracing used for quantitative analysis (E) Quantification of collagen fibers detected per ROI (F-H) Polar plots depicting collagen fiber orientation distributions for gel-only (F), NHLF-only (G), and NHLF + TGF-β (H) conditions (I) Fiber length distributions (J) Fiber straightness ratio (end-to-end distance ÷ curvilinear length) (K) Fiber width distributions. Data is presented as mean ± SEM. Statistical significance was determined by one-way ANOVA with post hoc testing. *p < 0.05, **p < 0.01, ***p < 0.001, ****p < 0.0001. Scale bars, 100 µm.

**Figure 2.**
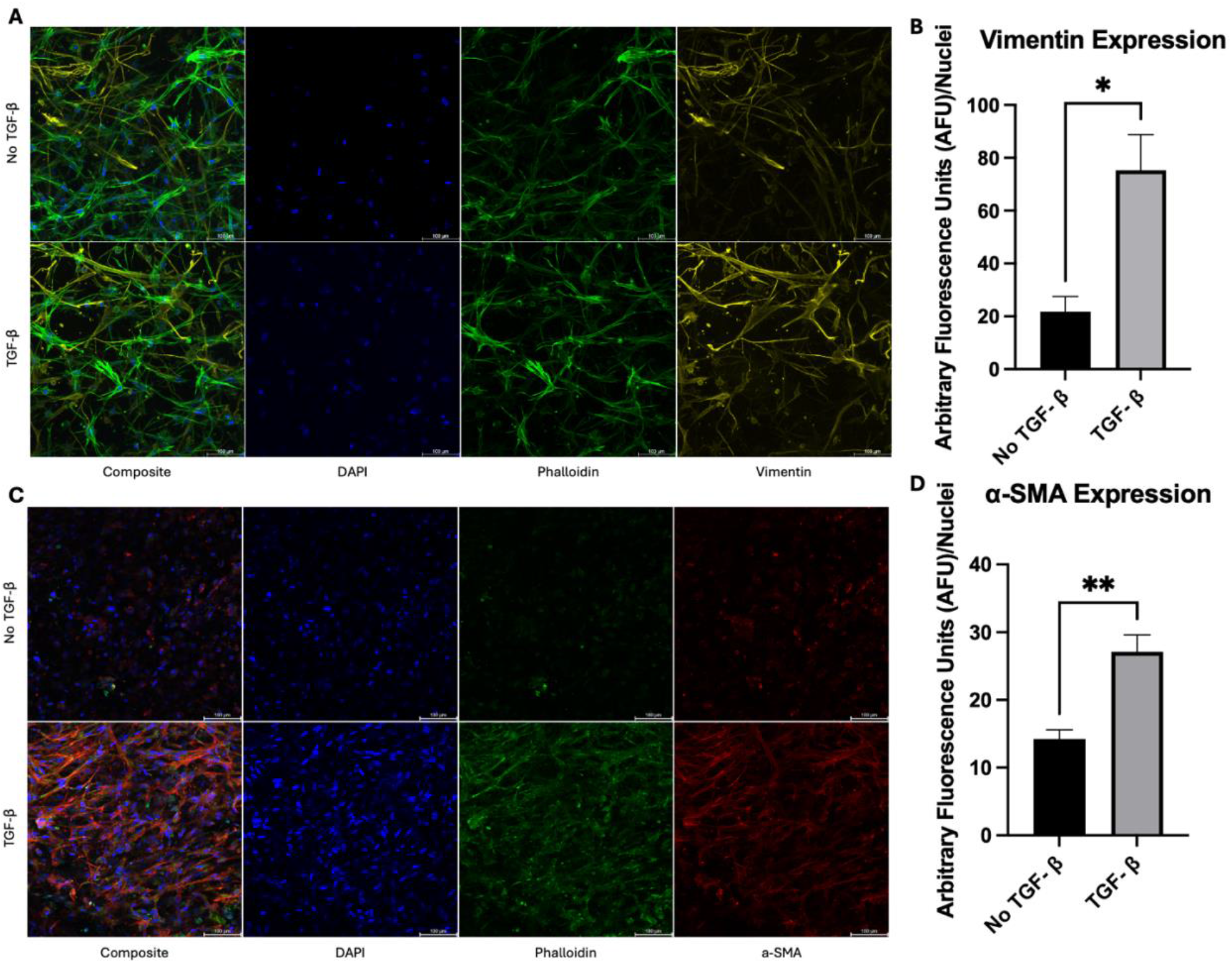
TGF-β enhances fibroblast activation in 3D culture. (A) Representative maximum intensity images of phalloidin (green)/Vimentin (yellow)/DAPI (blue) stain on NHLF for both no TGF-β and TGF- β groups, 10x (B) Vimentin expression per ROI (C) Representative maximum intensity images of phalloidin (green)/ α-SMA (red)/DAPI (blue) stain on NHLF (D) α-SMA expression per ROI, 20x Data are presented as mean ± SEM. Statistical significance was determined by one-way ANOVA with post hoc testing. *p < 0.05, **p < 0.01, ***p < 0.001, ****p < 0.0001. Scale bars, 100 µm.

**Figure 3.**
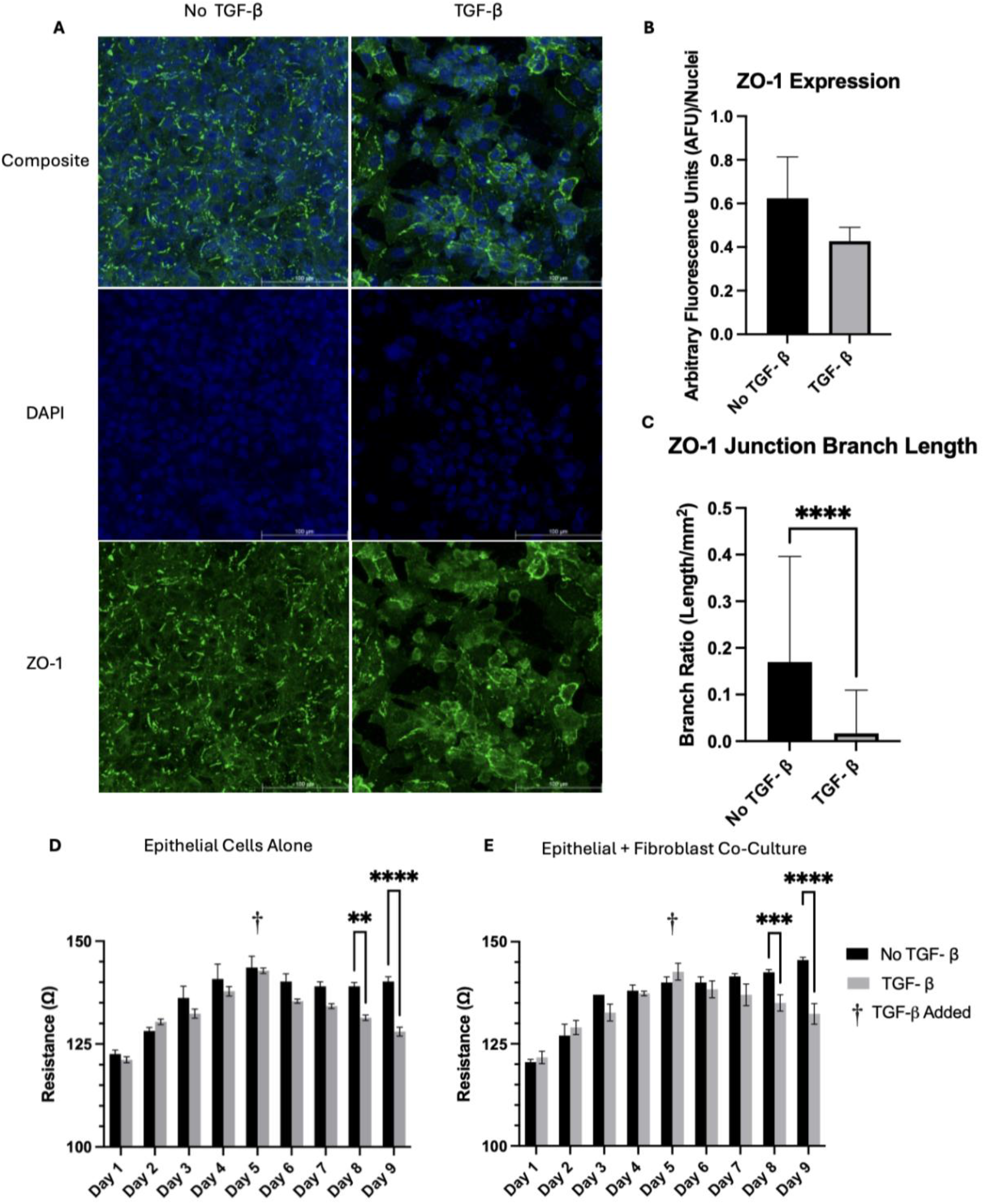
Exposure to TGF-β causes disruptions to epithelial barriers in the 3D co-culture model. (A) Representative images of ZO-1 (green)/ DAPI (blue) stain on A549s for both no TGF- β and TGF- β groups, 40x (B) ZO-1 expression per ROI (C) ZO-1 junction branch length per area of image (D) A549 barrier resistance over 9 day period. (E) Co-Culture of epithelial cells and fibroblast resistance over 9 day period. Data is presented as mean ± SEM. Statistical significance was determined by one-way ANOVA with post hoc testing. *p < 0.05, **p < 0.01, ***p < 0.001, ****p < 0.0001. Scale bars, 100 µm.

**Figure 4.**
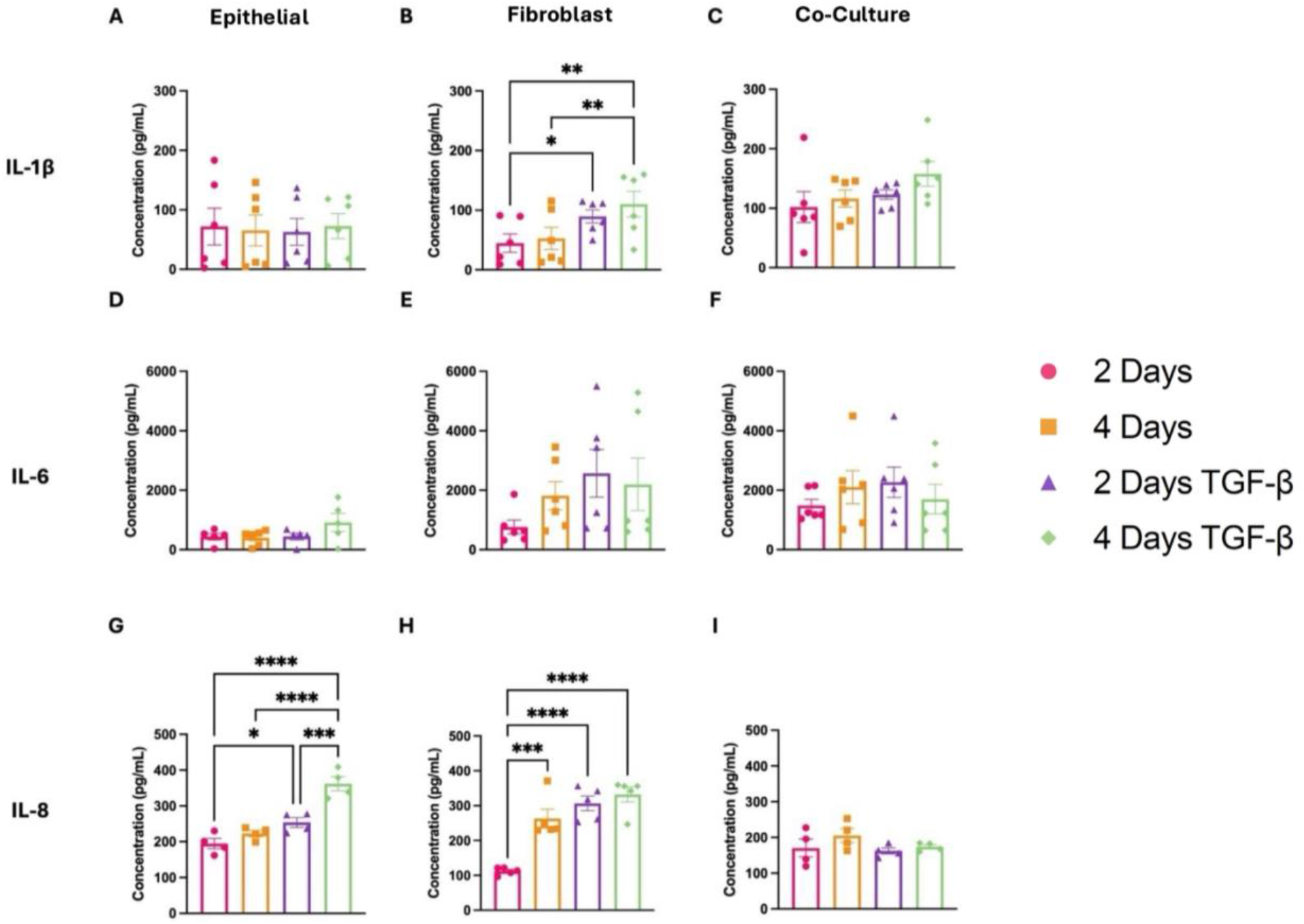
TGF-β Modulates IL-6, IL-1β, and IL-8 Expression in epithelial cells and fibroblasts. (A–C) IL-1β secretion over 4 days in NHLF monoculture, A549 monoculture, and co-culture with or without TGF-β. (D–F) IL-6 secretion over 4 days in NHLF monoculture, A549 monoculture, and co-culture with or without TGF-β. (G–I) IL-8 secretion over 4 days in NHLF monoculture, A549 monoculture, and co-culture with or without TGF-β. Statistical significance was determined by two-way ANOVA with post hoc testing. *p < 0.05, **p < 0.01, ***p < 0.001, ****p < 0.0001. Data are mean ± SEM; n ≥ 3 biological replicates per condition.

## 3. Results

### 3.1 TGF-β Induces Pronounced Collagen Remodeling and Collagen Fiber Alignment in 3D *in vitro* Lung Constructs

We observed distinct differences in collagen organization across experimental conditions. In acellular gels, the matrix appeared sparse and isotropic, with weak birefringence and predominantly fine, unbundled fibers lacking clear alignment (Figure 1A). Introduction of NHLFs led to a moderate increase in fiber density and brightness, along with localized bundling and partial alignment of collagen fibers (Figure 1B). These changes reflect fibroblast-mediated matrix remodeling, where cells exert mechanical tension on the surrounding ECM. In contrast, NHLF constructs treated with TGF-β exhibited markedly enhanced birefringence, characterized by thick, cable-like collagen bundles and a more cohesive directional orientation across larger regions (Figure 1C).

To quantify changes in collagen architecture across conditions, Picrosirius red-stained sections were analyzed using CT-FIRE (Figure 1D). The algorithm identified individual collagen fibers based on regional pixel intensity and morphology, extracting fiber-level parameters including length, width, straightness, and orientation. Overlays confirmed discrete fibrillar structures within regions of interest, allowing downstream quantification to reflect structural remodeling rather than bulk signal intensity. Acellular gel controls exhibited relatively few detectable fibers, consistent with a loosely organized, unstructured matrix. The inclusion of NHLFs resulted in a significant increase in the number of traceable fibers. TGF-β treatment further amplified this effect, driving a marked increase in fiber number (Figure 1E).

To quantify the degree of fiber alignment, a variety of different parameters of collagen orientation was assessed using CT-FIRE analyses. In acellular gels (Figure 1F), fiber angles were evenly distributed across the circular histogram, indicating an isotropic network with no dominant directionality. Constructs seeded with NHLFs (Figure 1G), exhibited a similar pattern with minor skewing toward vertical alignment, suggestive of localized tension and early fibroblast-mediated remodeling. In contrast, TGF-β–treated co-cultures (Figure 1H) demonstrated sharply peaked distributions centered near 0° and 180°, consistent with uniaxial alignment of collagen fibers and matrix anisotropy. Quantification of collagen length in gel-only constructs revealed that many of the detected fibers were short and fragmented, consistent with a randomly polymerized, non-remodeled hydrogel scaffold (Figure 1I). Upon seeding with NHLFs, the fiber length distribution shifted rightward. This effect was amplified under TGF-β stimulation, where constructs exhibited a significantly greater mean fiber length and an extended tail in the length distribution.

We also analyzed fiber straightness using the CT-FIRE–derived straightness ratio, which is defined as end-to-end distance divided by total curvilinear length for each fiber (Figure 1J). This metric quantifies fiber tautness, with values approaching 1 indicating straighter, more linearly extended fibers. Constructs composed without cells displayed the lowest straightness ratios, consistent with a relaxed, disorganized collagen network lacking cellular tension. The inclusion of NHLFs slightly increased fiber straightness, while TGF-β–treated cultures exhibited the highest straightness ratios, suggesting substantial fiber alignment and tension consistent with myofibroblast contractility. Finally, CT-FIRE outputs for the width of the detected fibers revealed a stepwise increase across experimental conditions, highlighting the progressive thickening of the ECM in response to cellular and biochemical cues (Figure 1K). Constructs composed of hydrogel alone exhibited the narrowest fibers, consistent with a baseline, unstructured matrix lacking cellular influence. Upon the introduction of NHLFs, fiber width increased significantly, reflecting fibroblast-driven bundling of collagen fibrils into thicker, more load-bearing structures. This trend was further amplified in TGF-β–treated cultures, where fiber width reached its highest levels, indicative of robust fibrillar aggregation and enhanced ECM remodeling. Taken together these findings indicate that activated NHLF activity drives a progressive transition from a sparse, isotropic network to a densely packed, highly aligned and tensioned collagen matrix that recapitulates key features of fibrotic ECM remodeling.

### 3.2 Fibroblast Activation is Amplified by TGF-β Treatment

TGF-β induced pronounced cytoskeletal remodeling within our hydrogel constructs. Compared with unstimulated gels, phalloidin staining on day 4 following TGF-β stimulation revealed increased formation of continuous F-actin stress fibers spanning the cell body (Figure. 2A). Vimentin staining likewise showed increased signal intensity and organization into thick, cable-like intermediate filament bundles extending across adjacent cells. Quantification demonstrated a significant increase in vimentin intensity per nucleus following TGF-β treatment relative to controls (Figure. 2B). TGF-β also increased α-SMA incorporation into filamentous stress fibers, with α-SMA co-localizing with phalloidin-labeled F-actin as continuous contractile bundles. In unstimulated gels, α-SMA signal was lower and largely diffuse. TGF-β treated hydrogels exhibited a denser, more aligned contractile network spanning neighboring cells. Per-cell analysis confirms a marked increase in α-SMA with TGF-β compared to control conditions (Figure 2D). These changes coincided with increased vimentin expression and extracellular matrix alignment and thickening (Figure 1).

### 3.3 TGF-β Disrupts Epithelial Barrier Integrity

Epithelial monolayers exposed to TGF-β exhibited marked junctional remodeling compared with untreated controls. ZO-1 immunofluorescence shifted from continuous, linear borders outlining uniformly sized cells to a more fragmented and discontinuous pattern in TGF-β–treated cultures, accompanied by enlarged and irregularly shaped cells (Figure 3A). Quantification of ZO-1 fluorescence intensity per nucleus showed a modest decrease with TGF-β treatment, whereas junction network analysis revealed a pronounced reduction in ZO-1 branch length per area (Figure 3B–C), indicating loss of tight-junction complexity. Consistent with these structural changes, TEER progressively declined in TGF-β–treated A549 monolayers relative to untreated controls over the 9-day period (Figure 3D), demonstrating impaired epithelial barrier function.

In the Transwell co-culture system, baseline TEER values were higher than in epithelial monocultures and increased during the pre-treatment period, reflecting barrier formation in the presence of fibroblasts (Figure 3E). Following addition of TGF-β to the basolateral compartment, co-cultures also displayed a significant reduction in TEER compared with untreated controls, indicating that pro-fibrotic signaling compromises barrier integrity even in the context of epithelial–fibroblast interactions.

### 3.4. TGF-β Modulates proinflammatory cytokine expression Across Cell Types

Cytokine analysis revealed distinct, cell type-dependent responses to TGF-β across epithelial, fibroblast and co-culture conditions. In epithelial monocultures, IL-1β levels remained low and unchanged over the 4 day period regardless of TGF-β exposure, indicating that epithelial cells are not a major source of IL-1β in this model (Figure 4C). IL-6 secretion was also low at baseline but showed a modest time-dependent increase by day 4 following TGF-β exposure, suggesting a limited epithelial contribution to the IL-6 milieu (Figure 4D). In contrast, IL-8 secretion was robustly induced by TGF-β exposure: levels increased between day 2 unstimulated and day 2 TGF-β–treated groups and rose further by day 4, consistent with strong, time-dependent chemokine production by epithelial cells (Figure 4G).

NHLF monocultures displayed a progressive and statistically significant increase in IL-1β secretion over the 4-day period, with a clear enhancement upon TGF-β stimulation, consistent with fibroblast activation and pro-inflammatory signaling (Figure 4B). NHLFs also produced substantially higher IL-6 than A549 cells at all time points; IL-6 increased from day 2 to day 4 and was further elevated by TGF-β, indicating that fibroblasts are the dominant source of IL-6 in this system (Figure 4E). IL-8 followed a similar pattern, with higher baseline levels than A549 cells and progressive up-regulation in response to TGF-β across all time points, underscoring the role of fibroblasts as key drivers of IL-8–mediated inflammation (Figure 4H).

In epithelial–fibroblast co-cultures, IL-1β levels were responsive to TGF-β-dependent, though increases were not statistically significant (Figure 4C). IL-6 levels in co-culture were intermediate between A549 and NHLF monocultures but retained a TGF-β-dependent increase, consistent with crosstalk that tempers fibroblast-derived IL-6 while maintaining a fibrotic cytokine signature (Figure 4F). Strikingly, IL-8 secretion in co-culture was reduced relative to both monoculture conditions and did not increase with TGF-β; instead, levels trended downward slightly, indicating that epithelial–mesenchymal crosstalk within the 3D system may actively suppress IL-8–driven inflammatory responses (Figure 4I).

## 4. Discussion

Idiopathic pulmonary fibrosis and related fibrotic lung diseases remain poorly understood in part because the cellular and mechanical interactions that drive disease progression are difficult to recapitulate in conventional culture systems. Two-dimensional monocultures models isolate individual features of fibrosis but cannot reproduce the reciprocal epithelial–mesenchymal signaling, matrix remodeling, and inflammatory dynamics that characterize progressive fibrotic injury in vivo. In this study, we established a 3D air–liquid interface co-culture model in which NHLFs are embedded in a collagen–hyaluronan hydrogel and layered beneath an epithelial compartment to mimic key features of the fibrotic lung microenvironment. Although the initiating insult in idiopathic pulmonary fibrosis remains unknown, transforming growth factor-β (TGF-β) is widely recognized as a key downstream mediator of the aberrant wound-healing response that drives fibrotic remodeling (Frangogiannis, 2020). TGF-β stimulation within this platform drove robust collagen remodeling, fibroblast activation, epithelial barrier disruption, and cytokine secretion in a cell type– and context-dependent manner. Picrosirius red staining of sectioned constructs combined with CT-FIRE analysis revealed progressive collagen reorganization from a sparse, isotropic network in cell-free gels to a densely packed, aligned, and thickened matrix in NHLF constructs. This was particularly evident after TGF-β treatment, consistent with matrix changes reported in fibrotic lung tissue and ECM-derived hydrogels. Following stimulation with TGF-β, NHLFs exhibited a myofibroblast-like phenotype with increased expression of vimentin, α-SMA, and F-actin stress fiber. At the epithelial level, basolateral TGF-β stimulation reduced TEER and shortened ZO-1 junction branch length, indicating impaired barrier integrity and altered tight junction organization. Finally, cytokine profiling showed that IL-1β, IL-6, and IL-8 were strongly induced in monocultures, whereas IL-8 secretion was dampened in epithelial–fibroblast co-cultures despite persistent IL-1β/IL-6, suggesting that bidirectional epithelial–mesenchymal crosstalk modulates the inflammatory milieu in a way not captured by monocultures alone.

Collagen remodeling is a central hallmark of fibrotic lung tissue, and multiple 3D hydrogel studies have demonstrated that matrix composition and stiffness regulate fibroblast phenotype and ECM production (Dabaghi et al., 2023; Guo et al., 2022; Phogat et al., 2023). Consistent with these findings, our results demonstrate that TGF-β stimulation drives quantitative increases in collagen fiber density, length, straightness, and width—features characteristic of an anisotropic matrix that mirrors fibrotic ECM architecture. This enhanced angular coherence reflects increased structural alignment and signifies a pathologic reorganization of the matrix that aligns with ECM architecture observed in IPF (Evanko et al., 2015; Singh et al., 2026).

Whereas many existing hydrogel studies focus primarily on fibroblast behavior or ECM mechanics in isolation, our platform extends these by allowing for evaluation of fibroblast-driven ECM remodeling and how this modulates epithelial barrier function and inflammatory signaling, providing a more integrated view of cellular-mechanical crosstalk during fibrosis. We previously demonstrated the capability of capturing ECM remodeling due to pro-inflammatory factors in the context of cancer (Nairon et al., 2022). Here we specifically focus on lung biology and an IPF-like state.

Several ALI models have shown that TGF-β can modulate epithelial tight junctions and barrier properties (Clerici and Matthay, 2003; Schilpp et al., 2021), although the direction of TEER change and junction remodeling appears to be context-dependent. In conductive airway epithelia, TGF-β1 redistributes tight junction proteins and increase permeability, while in some bronchial systems TGF-β or fibroblast co-culture increased TEER and junction protein expression (Schilpp et al., 2021). These studies highlight the importance of cell type, dose, and timing (Barron et al., 2024; Schilpp et al., 2021). In our model, TGF-β decreased TEER and reduced ZO-1 branch length, suggesting that pro-fibrotic signaling promotes barrier destabilization and junctional simplification, consistent with epithelial injury and fluidization observed in IPF-linked airway and distal epithelial cultures (Stancil et al., 2022). Our findings also complement epithelial–fibroblast ALI co-culture studies in which fibroblasts, particularly those derived from IPF patients, drive epithelial dysfunction, altered gene expression, and cytokine secretion (Barron et al., 2024; Prasad et al., 2014). By using normal fibroblasts in a fibrotic-like matrix and applying exogenous TGF-β, we model early or inducible fibroblast activation and show that the resulting ECM remodeling and cytokine milieu are sufficient to impair epithelial barrier properties, even in the absence of diseased primary cells.

The most striking finding in our cytokine data was the selective attenuation of IL-8 secretion in epithelial–fibroblast co-cultures relative to monocultures, despite sustained elevation of IL-1β and IL-6. This pattern suggests that direct epithelial–mesenchymal contact or paracrine signaling in this 3D ALI environment constrains IL-8–driven neutrophilic inflammation while preserving the IL-1β/IL-6 axis associated with fibrotic remodeling — a balance that may better approximate the chronic, low-grade inflammatory state of fibrotic lung tissue than acute monoculture responses. Dissecting the cellular sources of these mediators clarifies the picture further: fibroblasts were the dominant producers of IL-1β and IL-6, with TGF-β amplifying secretion over time, consistent with reports that activated fibroblasts sustain a pro-fibrotic cytokine environment (Ghonim et al., 2023; Seong et al., 2009; She et al., 2021). A549 epithelial cells contributed relatively little IL-1β or IL-6 but mounted a strong IL-8 response to TGF-β, in line with the established role of epithelial-derived IL-8 as a neutrophil chemoattractant and mediator of epithelial injury. That this IL-8 response was suppressed in co-culture — without a corresponding loss of IL-1β or IL-6 — points to a reshaping rather than a global dampening of the inflammatory milieu. Similar selective modulation of cytokine profiles has been reported in other co-culture systems where epithelial– mesenchymal or fibroblast–immune interactions constrain specific inflammatory pathways while reinforcing others (Barron et al., 2024; Herseth et al., 2008; Osei et al., 2020; Thiam et al., 2023), and our findings extend this pattern into a 3D fibrotic matrix context.

Our platform integrates several features that enhance physiological relevance compared with traditional 2D systems and simpler 3D gels. First, the collagen–hyaluronan hydrogel recapitulates critical ECM components present in fibrotic lungs, where altered collagen architecture and HA content influence fibroblast activation, TGF-β signaling, and tissue stiffness (Evanko et al., 2015; Guo et al., 2022; Herrera et al., 2018). Second, positioning the epithelial layer at an ALI on transwell inserts, with TGF-β applied to the basolateral chamber, reflects the polarized exposure of airway and distal epithelial cells to mesenchymal and vascular-derived cues during fibrosis progression. Third, direct epithelial–fibroblast co-culture enables reciprocal crosstalk via soluble factors and matrix remodeling, approximating the complex cellular interplay that drives fibroproliferation and barrier failure in idiopathic pulmonary fibrosis and other chronic lung diseases.

Although our work aligns with much of the previously published studies, there are some limitations to our culture system. We used immortalized A549 epithelial cells rather than primary distal airway or alveolar epithelium; although A549 cells are widely used and amenable to ALI culture, they only partially recapitulate the phenotype and injury responses of IPF-derived epithelial cells or iPSC-derived AT2-like cells used in more advanced models. Second, NHLFs were derived from a non-fibrotic donor. Future iterations of the model could study fibroblasts derived from explanted tissue of IPF patients. Although the collagen- and HA-rich hydrogel provides a complex ECM, future engineered ECM combinations and could be constructed to measure altered cross-linking, elastin loss, and disease-specific proteomic signatures. Additionally, our cytokine analyses focused on three canonical mediators and did not include broader panels of chemokines, growth factors, or matrix-modifying enzymes that regulate fibroblast–immune crosstalk and epithelial plasticity in IPF. Finally, the static Transwell configuration lacks dynamic mechanical stretch, interstitial flow, and immune cell trafficking, which may influence both matrix remodeling and cytokine gradients in vivo.

Future iterations of this platform could incorporate primary or iPSC-derived alveolar epithelial cells and fibroblasts from IPF patients to better capture disease-specific epithelial phenotypes and mesenchymal activation states. Integrating immune cells such as macrophages or neutrophils would enable direct study of fibroblast–immune–epithelial crosstalk and its contribution to persistent inflammation and ECM remodeling. Finally, adapting the system to microfluidic or lung-on-chip platforms could introduce controlled mechanical stretch and perfusion and realistic cytokine gradients, improving physiological relevance and enabling more precise modeling of spatial gradients and drug delivery. This engineered 3D lung platform provides a scalable system for mechanistic studies of fibrotic remodeling and for screening anti-fibrotic therapeutics.

## Conflict of Interest

The authors declare no competing interests.

## Data Availability

All data generated in this study are available from the corresponding author upon reasonable request.

## Notes

### Competing Interest Statement

The authors have declared no competing interest.

### Funding

- Ohio State University’s Center for Medical and Engineering Innovation

